# Demes: a standard format for demographic models

**DOI:** 10.1101/2022.05.31.494112

**Authors:** Graham Gower, Aaron P. Ragsdale, Gertjan Bisschop, Ryan N. Gutenkunst, Matthew Hartfield, Ekaterina Noskova, Stephan Schiffels, Travis J. Struck, Jerome Kelleher, Kevin R. Thornton

## Abstract

Understanding the demographic history of populations is a key goal in population genetics, and with improving methods and data, ever more complex models are being proposed and tested. Demographic models of current interest typically consist of a set of discrete populations, their sizes and growth rates, and continuous and pulse migrations between those populations over a number of epochs, which can require dozens of parameters to fully describe. There is currently no standard format to define such models, significantly hampering progress in the field. In particular, the important task of translating the model descriptions in published work into input suitable for population genetic simulators is labor intensive and error prone. We propose the Demes data model and file format, built on widely used technologies, to alleviate these issues. Demes provides a well-defined and unambiguous model of populations and their properties that is straightforward to implement in software, and a text file format that is designed for simplicity and clarity. We provide thoroughly tested implementations of Demes parsers in multiple languages including Python and C, and showcase initial support in several simulators and inference methods. An introduction to the file format and a detailed specification are available at:https://popsim-consortium.github.io/demes-spec-docs/.

## Introduction

The ever-increasing amount of genetic sequencing data from genetically and geographically diverse species and populations has allowed us to infer complex demography and study life history at fine scales. An integral component to such population genetics studies is simulation. Software to either simulate whole genome sequences (Thornton, 2014, 2019, Staab *et al*., 2015, Baumdicker *et al*., 2022, Kelleher *et al*., 2016, Haller and Messer, 2019) or informative summary statistics of diversity (Gutenkunst *et al*., 2009, Kamm *et al*., 2017, Jouganous *et al*., 2017) have enabled the increasing complexity of genomic studies, with several software packages capable of handling large sample sizes, many interacting populations, and deviations from panmictic random-mating assumptions. This ability to infer and simulate such complex demographic scenarios, however, has highlighted a major shortcoming in community standards: the fragmented landscape of different ways to describe demographic models makes it difficult to compare inferences made by different methods and to reliably simulate from previously inferred models. Inference results are typically reported in publications via a combination of visual depiction, a list of key parameters in tabular form and a discussion within the text. Unfortunately these descriptions are often ambiguous, and implementing the precise model inferred for later simulation is at best tedious and error prone (Adrion *et al*., 2020, Ragsdale *et al*., 2020), and occasionally impossible because of missing information. Simulation is a core tool in population genetics, and many methods have been developed over the past three decades (Carvajal-Rodríguez, 2008, Liu *et al*., 2008, Arenas, 2012, Yuan *et al*., 2012, Hoban *et al*., 2012). Simulations are based on highly idealized population models, and one of the key uses of inferred demographic histories is to make simulations more realistic. Simulation methods take three broad approaches to specifying the demographic model to simulate, using either a command line interface (e.g., Hudson, 2002, Hernandez, 2008, Kern and Schrider, 2016), a custom input file format (e.g., Guillaume and rougemont, 2006, Excoffier and Foll, 2011, Shlyakhter *et al*., 2014), or an Application Programming Interface (API) to allow models to be defined programmatically (e.g., Thornton, 2014, Hernandez and Uricchio, 2015, Kelleher *et al*., 2016, Becheler *et al*., 2019, Haller and Messer, 2019, Thornton, 2019, Baumdicker *et al*., 2022). Command line interfaces are a concise way of expressing demographic models, and the syntax defined by `ms` (Hudson, 2002) is used by several simulators (e.g., Ewing and Hermisson, 2010, Chen *et al*., 2009, Staab *et al*., 2015). However, this conciseness means that models of even intermediate complexity are difficult for humans to understand, making errors likely. APIs are more verbose, but require a substantial time investment to learn, and as they are tied to a specific tool this knowledge is not portable to other simulators. Like APIs, input parameter file formats for simulators allow the model specification to be less terse and allow for documentation in the form of comments. Several graphical user interfaces and visualization methods have been developed, which greatly facilitate interpretation (Mailund *et al*., 2005, Antao *et al*., 2007, Parreira *et al*., 2009, Ewing and Hermisson, 2010, Parobek *et al*., 2017, Zhou *et al*., 2018). However, these methods currently have little traction as they are all either directly coupled to an internal simulation method or to the syntax of a specific simulator. There is currently no way in which demographic models inferred by different packages can be simulated or visualized by downstream software.

Here we present “Demes”, a data model and file format specification for complex demographic models developed by the PopSim Consortium (Adrion *et al*., 2020). The Demes data model precisely defines the sizes and relationships of populations, and it provides a way to explicitly encode the information relevant to demography while avoiding repetition. This data model is implemented in the widely used YAML format (Ben-Kiki *et al*., 2009), which is a data serialization language that provides a good balance between human and machine readability. The specification precisely defines the required behavior of implementations, ensuring that there is no ambiguity of interpretation, and includes both a reference implementation and an extensive suite of test examples and their expected output. The initial software ecosystem includes high-quality Python and C parser implementations, as well as utilities for verification and visualization of Demes models, and has been implemented in several popular inference and simulation methods (Table 1). We hope that this data model and file format will be widely adopted by the community, such that users can expect to simulate directly from inferred models with little to no programming effort.

**Table 1.**
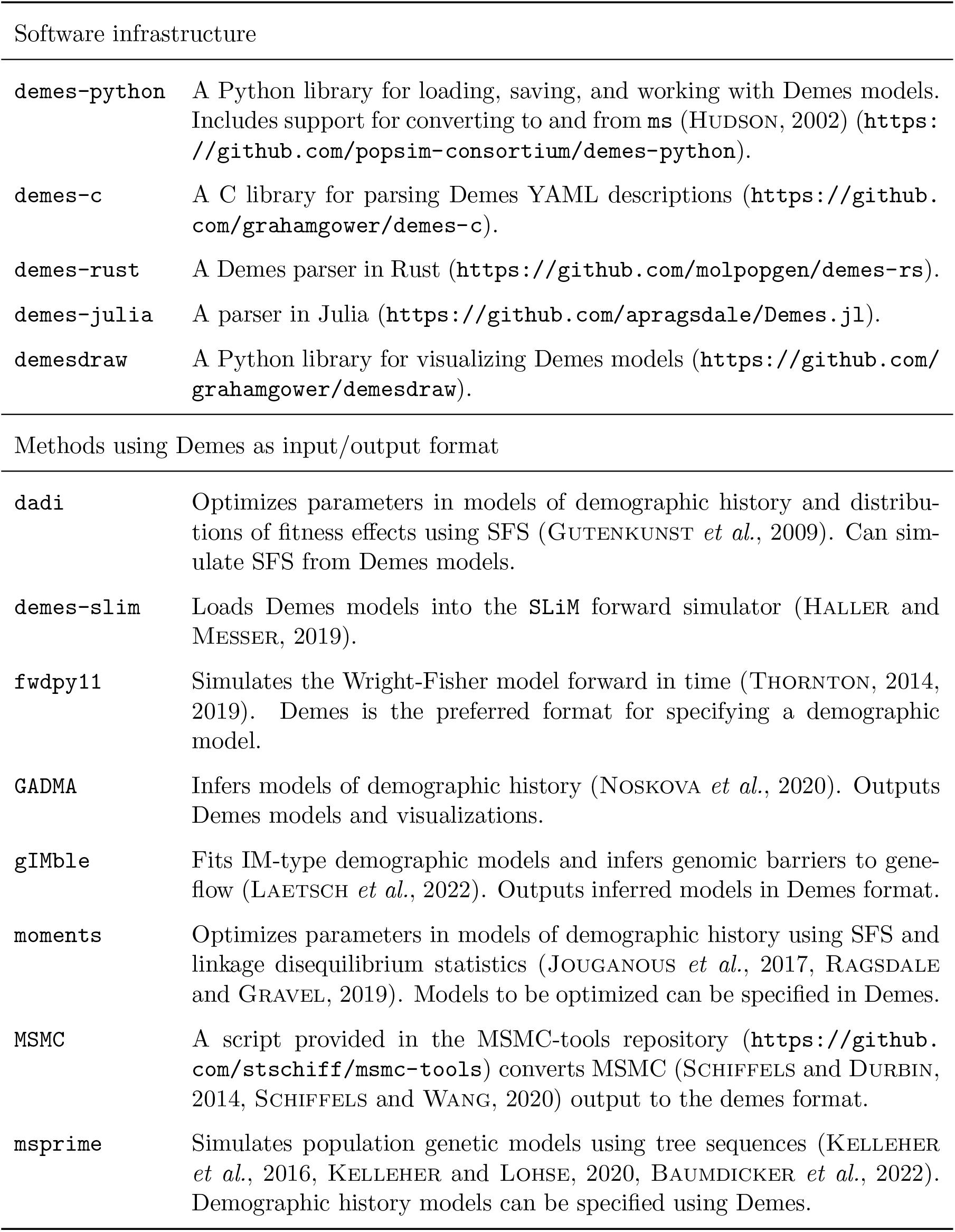
Software support for Demes. We have included software infrastructure developed for working with Demes models (such as parsing, validation, and visualization) as well as downstream software that implement the specification, at the time of writing.

## Demes

The design of Demes is a balance between two partially competing requirements: that (a) models should be easy for humans to understand and manipulate; and (b) software processing Demes models should be provided with an unambiguous representation that is straightforward to process. For efficiency of understanding and avoidance of model specification error, we require a data representation without redundancy (i.e., repetition of values). However, for the simplicity of software working with the Demes model (and the avoidance of programming error, or divergence in interpretations of the specification) it is preferable to have an explicit representation, in which all relevant values are readily available. Thus, Demes is composed of three entities: the Human Data Model (HDM) designed for human readability; the Machine Data Model (MDM) designed for programmatic input and processing; and the parser, which is responsible for transforming the former into the latter.

Here we provide a brief overview of the population genetics models that Demes supports and the components of the Demes infrastructure. Complete technical details of the MDM and HDM, and the responsibilities of the parser are provided in the online Demes specification (https://popsim-consortium.github.io/demes-spec-docs/). This specification rigorously defines the data model, fully describing the entities and their relationships, and the required behavior of implementations. Since the online specification is definitive, we will not recapitulate the details here, but instead focus on the high level properties of the model and the rationale behind key design decisions.

### Population genetics model

For inference and simulation software to meaningfully interoperate there must be a shared understanding of what a demographic model *is*. Population genetics is a large field, and rather than attempting to capture all possible within- and between-population processes, we have instead adopted a pragmatic approach of identifying a common set of assumptions shared by many methods. We outline the processes and assumptions briefly here and in Appendix A1.

Demographic models consist of one or more populations (or “demes”) defined by their size histories and the time intervals of their existence. Individuals can move between populations based on their ancestor-descendant relationships or by continuous or discrete migration events. Within a population, we assume Wright-Fisher dynamics (see Appendix A1.3 for more precise details). As described in the Scope of the Specification section below, the demographic model does not, as a deliberate simplification and separation of duties, include any information about genome biology or selection.

These basic assumptions of discrete Wright-Fisher populations connected by instantaneous or continuous migrations are shared by many inference methods (e.g., Gutenkunst *et al*., 2009, Li and Durbin, 2011, Gravel, 2012, Schiffels and Durbin, 2014, Kamm *et al*., 2017, Jouganous *et al*., 2017, Ragsdale and Gravel, 2019, Excoffier *et al*., 2021), and forwards- and backwards-time simulators (e.g., Hudson, 2002, Gutenkunst *et al*., 2009, Excoffier and Foll, 2011, Kelleher *et al*., 2016, Jouganous *et al*., 2017, Haller and Messer, 2019, Thornton, 2019). Demes therefore serves as “middleware” between inference methods and simulation software, capturing these common assumptions.

It is important to note that the goal of describing the basic population processes precisely is not to be proscriptive about what methods may or may not use the specification, but so that we can be clear on what situations we can expect methods to agree exactly. Arbitrary population processes —for example, within-deme continuous spatial structure (Wright, 1943, Barton*et al*., 2002, 2010, Ringbauer *et al*., 2017, Battey *et al*., 2020)—may be layered on top of this basic description, but as dynamics diverge from the core assumptions, then of course we can expect results to differ accordingly.

### Human Data Model

The Demes Human Data Model (HDM) is focused on efficient human understanding and avoiding errors. We have adopted the widely used YAML format (Ben-kiki *et al*., 2009) as the primary interface for writing and interchanging demographic models (see Appendix A2 for rationale). Demographic models provide information about global features of the model (such as time units and generation times), populations (as “demes”) and their existence intervals (as “epochs”), and gene flow between populations (as continuous “migrations” or instantaneous “pulse” events). Fig 1 shows an example isolation-with-migration model in HDM format.

**Figure 1:**
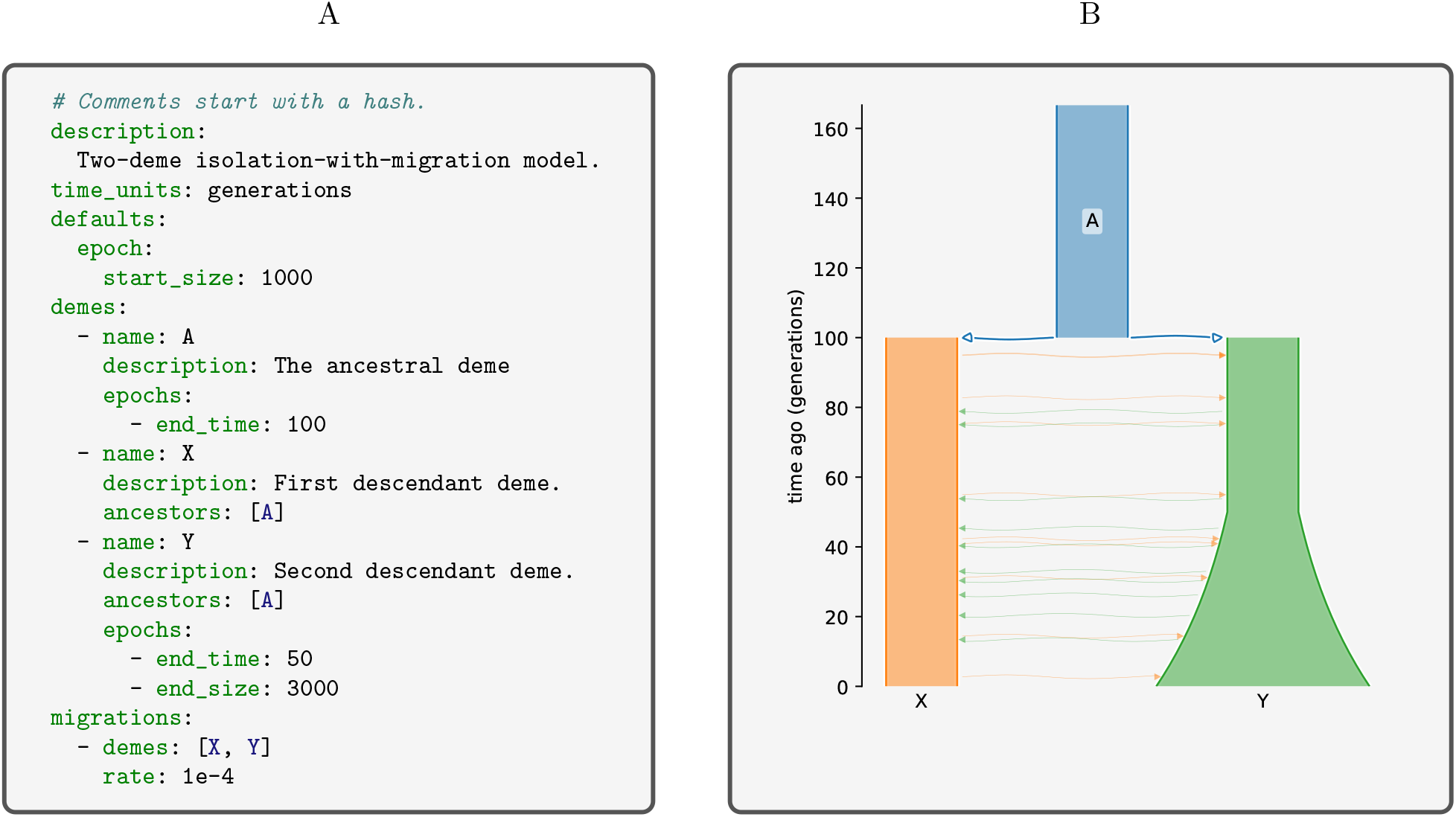
Example isolation-with-migration Demes model. (A) The Human Data Model representation expressed using YAML. (B) A visual representation of the model using `demesdraw`. The same model in the Machine Data Model form is provided in Figure A1.

Structurally, the HDM encourages human understanding by avoiding redundancy in the description where possible and by providing a mechanism for specifying default values that are inherited hierarchically. For values that repeat across fields, the “defaults” mechanism may be used to implicitly assign default values to fields of the given type. A default is superseded by an explicitly provided value if given. Size values are inherited naturally following the progression of time. For example, if an epoch `start size` is not provided (either directly, or via a defaults section), it is assumed to be equal to the `end size` of the previous epoch. This also means that the first epoch of each population must specify the initial size (or it must be provided in a defaults section).

Avoiding redundancy in this way reduces the cognitive load on readers, by highlighting necessary parameters which may be otherwise be obscured. It is not necessary—or indeed recommended— that all models are expressed in a maximally concise form, and we wholeheartedly endorse the explicit statement of parameters where it increases model legibility.

### Parsers

While the HDM is designed for human readability and conciseness, the underlying data model suitable for software implementation (the Machine Data Model, or MDM) is redundant and exhaustive. Translation from the HDM to the MDM requires resolving hierarchically-defined default values and verifying relationships between populations and the validity of specified parameter values. Because this translation and validation requires significant programming effort, we define a standard software entity as part of the specification to perform this task (the parser), which is intended to be shared by programs that support Demes as input. The Demes specification precisely defines the required behavior of parsers, and we provide a reference implementation written in Python to resolve any potential ambiguities, as well as an extensive test suite of examples and the expected outputs. In addition, we have high-quality parser implementations in the Python, C, Rust, and Julia languages (Table 1) providing a solid foundation for the software ecosystem. By maintaining high-quality Demes parsers available as libraries, we ensure consistency across simulation and inference software. Having common parsers also benefits users by providing consistent and informative error messages for missing values or issues in formatting.

### Scope of the specification

A primary design goal of Demes is to provide a means of unambiguously communicating the results of demographic model inferences to population genetic simulators. Since demography is defined in terms of groups of individuals and these groupings are influenced by genetics, it is difficult to find a simple definition that separates the two. Thus, we have attempted to be pragmatic, limiting the features that we include in Demes to those that are in practise regarded as part of a demographic model.

The model is therefore limited to features that we can expect many different demographic inference and simulation methods to share. The specification only describes demographic features at the population level. Features of genome biology are out of scope, including mutation and recombination rates, genome annotations, ploidy, and so on. Selection and dominance models are absent, as discussed in Appendix A1. It is important to note, however, that Demes may be used in applications that include additional population genetic processes outside of what is explicitly modeled in the specification, such as interpreting population sizes as carrying capacities, implementations of hard selection, or layering more complicated mating or spatial structure. The Demes specification is intended to provide a basic model that can be elaborated on where necessary.

Demes is not a standard population genetic simulation specification, although it could be *part* of one. Since the standard is based on JSON, and JSON documents can be arbitrarily nested, we can imagine a simple specification of genome features such as mutation and recombination rates in which the demography is defined by an embedded Demes specification. Features of the simulation specification (such as defining the time and location of samples) can then *refer to* the Demes model. This design, in which we embed the demographic model *within* a larger specification rather than adding arbitrary and unrelated complexities *to* the demography is an essential simplification and separation of duties.

The Demes specification is static by design—we wish to unambiguously describe a demographic model with a concrete set of parameters. This simplicity means that we cannot directly specify parameter distributions or estimated confidence intervals for those parameters. While it is not difficult to imagine extending the specification in ways that would allow this, it is not clear that the benefits are worth the greatly increased parser complexity (see Appendix A3).

## Example: an isolation-with-migration model

In Figure 1 we provide an example isolation-with-migration model. Models typically start with a concise description, followed by the mandatory `time units` field. This model uses the `defaults` section to provide a default `start size` of 1000 individuals for each epoch of each deme. There are three demes in the model, an ancestral deme named “A” which exists arbitrarily far back into the past then ceases to exist at 100 generations ago, and demes “X” and “Y” that derive their ancestry from A when it goes extinct. Demes A and X have only one epoch, in which the population sizes are constant, whereas deme Y has two epochs. Deme Y’s second epoch has a different `end size` than its `start size`, which indicates the size grows exponentially from 1000 individuals at 50 generations ago to 3000 individuals at time 0 (the present). The migration section lists one migration stanza, between demes X and Y. This migration stanza doesn’t indicate a source or destination deme, so the migration is symmetric. No migration times are specified, so migrations occur continuously at the given rate during the time interval over which both demes exist (from 100 generations ago until the present). We do not attempt a detailed explanation of all Demes features here, and readers are instead directed to the tutorial and detailed specification in the online documentation (https://popsim-consortium.github.io/demes-spec-docs/).

## Application: simulation using Demes

Here, we highlight the interaction between Demes and other software, including simulation and model illustration tools. Demes allows us to specify a demographic model which can be used as the input for a growing number of simulation packages (Table 1). We implemented the human two-population demographic model from Tennessen *et al*. (2012) inferred from European and African-American sequencing data. This model (shown in Demes format in Figure A2) is parameterized by an ancestral population with an ancient growth, divergence into “AFR” and “EUR” that each have multiple-epoch size histories, and multiple epochs of continuous migration between the two branches (illustrated using `demesdraw` in Figure 2A). The large final sizes (≈ 500, 000 individuals each) are one to three orders of magnitude larger than ancestral population sizes, reflecting the recent explosive population size increase in humans.

**Figure 2:**
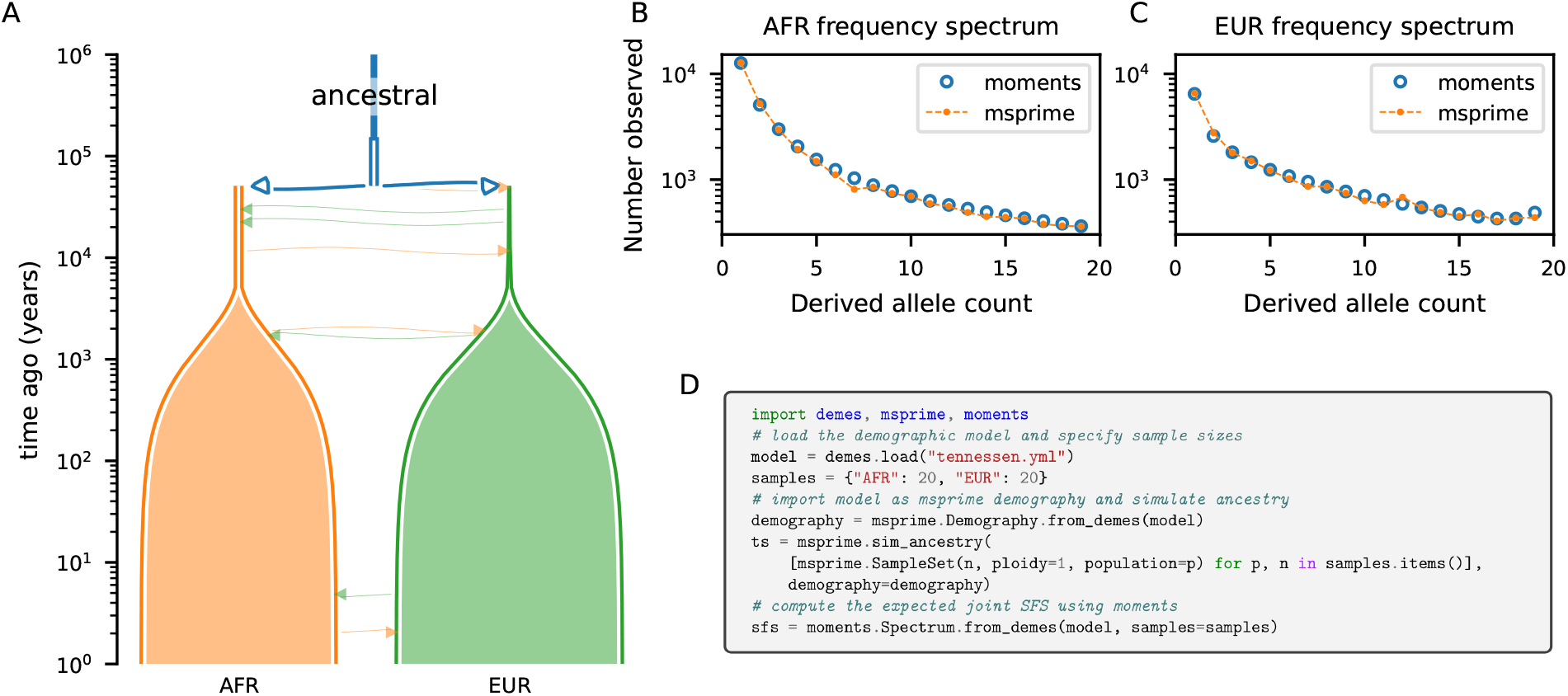
Illustration and simulation using Demes. (A) Using an inferred demographic model from Tennessen *et al*. (2012) specified as a YAML file in Demes format (Figure A2), we used `demesdraw` to visualize the demographic model (note the recent exponential growth resulting in present-day population sizes that greatly exceed those in the past). We then used `msprime` to simulate genomic data for 20 genome copies sampled from the two contemporary populations, and we used `moments` to compute the expected joint site-frequency spectrum for the same sample sizes (Figure A3). (B, C) We compared the single-population SFS in each population, showing agreement between the simulation methods. (D) Python code snippets of the interactions between `demes` and the simulation software. An extended script to compute the SFS shown in (B) and (C) is given in Figure A3.

We used this model to simulate 20 haploid genome copies from EUR and AFR at time zero (i.e., present day) to obtain the joint site-frequency spectrum (SFS), a summary of observed allele frequencies widely used in evolutionary inference (Bustamante *et al*., 2001, Gutenkunst *et al*., 2009, Tennessen *et al*., 2012, Jouganous *et al*., 2017, Kamm *et al*., 2017, Kim *et al*., 2017). The Demes model (Figures 2A and A2) was provided as the input demography to `msprime` (Baumdicker *et al*., 2022) to simulate a large recombining region under the mutation rate assumed in Tennessen *et al*. (2012), and we computed the observed SFS using `tskit` (Ralph *et al*., 2020). Using the same Demes model as input to `moments` (Jouganous *et al*., 2017), we computed the expectation of the joint SFS and compared to the `msprime` simulated data (Figure 2B,C). Figure 2D shows the code required to run the simulations in `msprime` and `moments`, and demonstrates that precisely the same input model, without modification, was provided to both packages. Such interoperability is a major gain for researchers, which we hope will become the expected norm as more packages adopt the Demes format.

## Discussion

Stable and healthy software ecosystems require standard interchange formats, allowing for the development of high-quality and long-lasting tools that produce and consume the standard. Demographic models are a key part of population genetics research, and to date the transfer of inferred models to downstream simulations has been *ad-hoc*, and conversions between the many different ways of expressing such models is both labor intensive and error-prone. The proposed Demes standard is an attempt to bridge this gap between inference and simulation, and also to provide the foundations for a sustainable ecosystem of tools built around this data model. Table 1 shows some initial infrastructure that we have built as part of developing Demes, but many other useful tools can be envisaged that produce, consume, or transform this format.

Reproducibility is a significant problem throughout the sciences (Baker, 2016), and various measures have been proposed to increase the likelihood of researchers being able to replicate results in the literature (MunafÓ *et al*., 2017). The most basic requirement for reproducibility is that we must be able to state precisely what the result in question *is*. The lack of standardization in how complex demographic models are communicated today, and the lack of precision in the published model descriptions means that it is difficult to replicate analyses, or reproduce those models for later simulation. Thus, we hope that the Demes standard introduced here will be widely adopted by simulation and inference methods and be used for reporting results in publications, either as supplemental material or uploaded to a data repository.

## Acknowledgments

We would like to thank the editor and reviewers for helpful comments that have significantly improved this manuscript. Graham Gower was supported by a Villum Fonden Young Investigator award to Fernando Racimo (project no. 00025300). Ryan Gutenkunst and Travis Struck were supported by the National Institute of General Medical Sciences of the National Institutes of Health (R01GM127348 to RNG). Matthew Hartfield is supported by a NERC Independent Research Fellowship (NE/R015686/1). Jerome Kelleher is supported by the Robertson Foundation. Stephan Schiffels was supported by funding from the European Research Council (ERC) under the European Union’s Horizon 2020 research and innovation programme (grant agreement No 851511). Gertjan Bisschop was supported by funding from the ERC(ModelGenomLand, 757648).

## Appendix

The Demes specification is a formal data model for describing the properties of populations over time, along with some metadata and provenance information. The data model is based on the ubiquitous JSON (Bray, 2017) standard, and formally defined using JSON Schema (Wright *et al*., 2020). Along with the schema, full technical details of the of the model are provided in the online specification document (https://popsim-consortium.github.io/demes-spec-docs/).

### A1 Population genetics model details

In Demes, demographic models consist of one or more interacting populations, or “demes”, understood to be a collection of individuals that can be conveniently modeled using a defined set of rules and parameters (Gilmour and Gregor, 1939, Gilmour and Heslop-Harrison, 1955). To avoid confusion with the name of the specification itself we will use the term “population” in this discussion, with the understanding that the terms are interchangeable. A population is defined as some collection of individuals that exists for some period of time, and has a well-defined size (i.e., number of individuals) during that time period. Individuals can move between populations either according to their ancestor-descendant relationships or through processes involving migrations. Few other properties of the populations are specified in the model: we are concerned primarily with defining the populations, their sizes, and the movement of individuals between those populations.

#### A1.1 Time units

Population and event times are written as units in the past, so that time zero corresponds to the final generation or “now”, and event times in the past are values greater than zero with larger values corresponding to times in the more distant past. By having time values increase into the past, we avoid the need to choose an arbitrary point in history as “time zero”. A natural specification for time units is in generations, although other time units are permitted, such as years, accompanied by the generation time so that downstream software may convert times into generations as required. There must be at least one population with an infinite `start time`. An infinite start time may be interpreted differently depending on the simulator. In a coalescent setting, there is no upper bound for the coalescent time of lineages in this population. In a forwards-time setting, the interval of time between infinity and the oldest non-infinite model time (i.e. the “first event”) is approximated by the simulator’s burn-in phase—detailed guidance is provided in the online specification.

#### A1.1 Sizes and epochs

Population sizes are given as numbers of individuals, and details such as ploidy levels are considered external to the model. We therefore focus on the number of individuals as opposed to the number of genome copies. Sizes and mating system details are specified for each population within population-specific epochs. Epochs are contiguous time intervals that define the existence interval of the population. Each epoch specifies the population size over that interval, which can be a constant value or a function defined by start and end sizes that must remain positive. Only exponential population size changes are currently supported, but other functions may be added to the specification over time.

#### A1.1 Population dynamics

Within a population, we assume that allele frequency dynamics can be described by the Wright-Fisher model. Briefly, generations are non-overlapping (all parents reproduce and die simultaneously), and for allele *i* currently at frequency *p*_*i*_, its frequency in the next generation (at birth) is expected to be 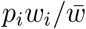, where *w*_*i*_ and 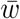 are the marginal and mean fitnesses, respectively, properly weighted according to ancestry proportions. In this framework, a forward-time simulation of finite populations is equivalent to multinomial sampling of allele frequencies each generation (Bürger (2000, pp 29-31), Crow and Kimura (1970, pp 179-181)), and a backwards-time (coalescent) simulation follows the approximations described in Tajima (1983), Hudson (1983) and Wakeley (2008, chapter 3). Further, this model assumes “soft” selection (Christiansen, 1975), meaning that the dynamics of population sizes changes are independent of the details of individual fitnesses. As such, this model excludes scenarios such as “hard selection,” in which population sizes are dependent on a population’s mean fitness, or stochastic fluctuations in population size, such as interpreting population sizes as carrying capacities. Many forwards and backwards time simulators currently implement this model (e.g., Hudson, 2002, Gutenkunst *et al*., 2009, Excoffier and Foll, 2011, Kelleher *et al*., 2016, Jouganous *et al*., 2017, Haller and Messer, 2019, Thornton, 2019).

#### A1.1 Selfing and cloning

Each population has an assigned selfing rate and cloning rate, where each defines the probability that offspring are generated from one generation to the next by either self-fertilization or cloning of an individual. More specifically, for a given epoch within a population denote the clonal rate by *σ* and the selfing rate by *S. S* and *σ* can take any value between zero and one and can sum to more than one. Each generation a proportion of offspring *σ* are expected to be generated through clonal reproduction, while 1 − *σ* are expected to arise through sexual reproduction. Within the sexually-reproduced offspring, a proportion *S* are born via self-fertilization while the rest have parents drawn at random from the previous generation. Depending on the simulator, this random drawing of parent may occur either with or without replacement. When drawing occurs with replacement, a small amount of “residual” selfing is expected, so that the realized selfing probability is (1 − *σ*)(*S* + (1 − *S*)*/N*) instead of (1 − *σ*)*S* (so that even with *σ* = 0 and *S* = 0, selfing may still occur with probability 1*/N*), although this effect is negligible in large populations (Nordborg and Donnelly, 1997).

By allowing the definition of selfing and cloning probabilities, we allow many standard models to be defined. However, by parameterizing selfing and cloning as we have, we assume that these properties of populations can be specified independently from the genetics. In other words, mutations that cause selfing probabilities to fluctuate within an epoch are not considered. More details of the mathematical properties of selfing and cloning rates in a coalescent context can be found in Nordborg and Donnelly (1997), Hartfield *et al*. (2016).

#### A1.5 Relationships between populations

A population may have one or more ancestors, which are other populations that exist at the population’s start time. If one ancestor is specified, the first generation is constructed by randomly sampling parents from the ancestral population to contribute to offspring in the newly generated population. If more than one ancestor is specified, the proportions of ancestry from each contributing population must be provided, and those proportions must sum to one. In this case, parents are chosen randomly from each ancestral population with probability given by those proportions.

Individuals in a population may have parents from a different population through migrations. These can be defined as continuous migration rates over time intervals for which populations co-exist or through instantaneous (or pulse) migration events at a given time. Continuous migration rates are defined as the probability that parents in the “destination” population are chosen from the “source” population. On the other hand, pulse migration events specify the instantaneous replacement of a given fraction of individuals in a destination population by individuals with parents from a source population.

### A2 Rationale for YAML

We have adopted the widely used YAML format (Ben-Kiki *et al*., 2009) as the recommended means of interchanging Demes models (e.g., Figures 1 and A2). YAML is a data serialization language with an emphasis on simplicity and which interoperates well with JSON (indeed, YAML 1.2 is a superset of JSON). We chose YAML over JSON because although JSON is an excellent format for data interchange, it is ill-suited for human understanding and manipulation. We also considered other declarative data exchange formats such as TOML, but chose YAML because of its equivalence with JSON, popularity, and good software support. Since the Demes data model is defined in JSON Schema, however, there is no formal dependency on YAML and implementations may choose to use JSON directly if they wish (e.g., for greater efficiency).

### A3 Rationale for static models

The Demes specification is designed to describe demographic models defined by a fixed set of model parameters. As described in the main text, it does not include information about estimated confidence intervals or the joint distribution of parameter values. In this section we describe the rationale for this design decision.

The parameters of demographic models are typically tightly coupled, and cases in which distributions for different parameters can be simply described are rare. In this situation, the simplest way to describe an estimated distribution is to list a large number of samples from the posterior. While writing out a large number of Demes models in YAML format may seem inefficient, it can in fact be a compact way to describe these distributions. For example, consider a one-population model with piecewise-constant sizes over 20 epochs which has ∼ 40 free parameters: the `start size` and `end time` values for each epoch. If we sample 50,000 models from the posterior distribution, the resulting multi-document YAML file is 45 MiB. This format compresses down to 8.4 MiB when gzipped or 6.2 MiB when compressed with LZMA2, which is on par with an equivalent binary representation of the free parameters (40 × 50000 × 4 bytes ≈ 7.6 MiB).

Similarly, one might be interested in running simulations in which the demographic model parameters are drawn from a distribution, e.g., in ABC inference (Beaumont *et al*., 2002). Other inference procedures based on optimizing a loss function (Gutenkunst *et al*., 2009, Kamm *et al*., 2017, Jouganous *et al*., 2017, Ragsdale and Gravel, 2019, Excoffier *et al*., 2021) need users to specify parameter bounds, and possibly non-linear or conditional constraints between parameters. Indeed, the choice of how to parameterize a model could be important for some inference methods (e.g. absolute times versus relative times between events).

Implementing the many distributions of interest and supporting a general way to describe a model’s free parameters would greatly increase the complexity of parsers, with relatively limited benefit to most users. It is unlikely that Demes could be made sufficiently flexible without implementing many features of general-purpose programming languages, such as variables, arithmetic, and flow control. Such use cases are therefore better served by writing model-generating functions in an existing programming language, for example using the Demes Python API (e.g., as implemented in `moments` (Jouganous *et al*., 2017, Ragsdale and Gravel, 2019)). As an intriguing possibility for developments in this direction, there exist many templating solutions for YAML and JSON that are specifically designed for extending static data in arbitrarily complex ways (e.g., `YTT`, `Jsonnet`, `CUE`, and `Dhall`).

### A4 Extended data and figures

**Figure A1:**
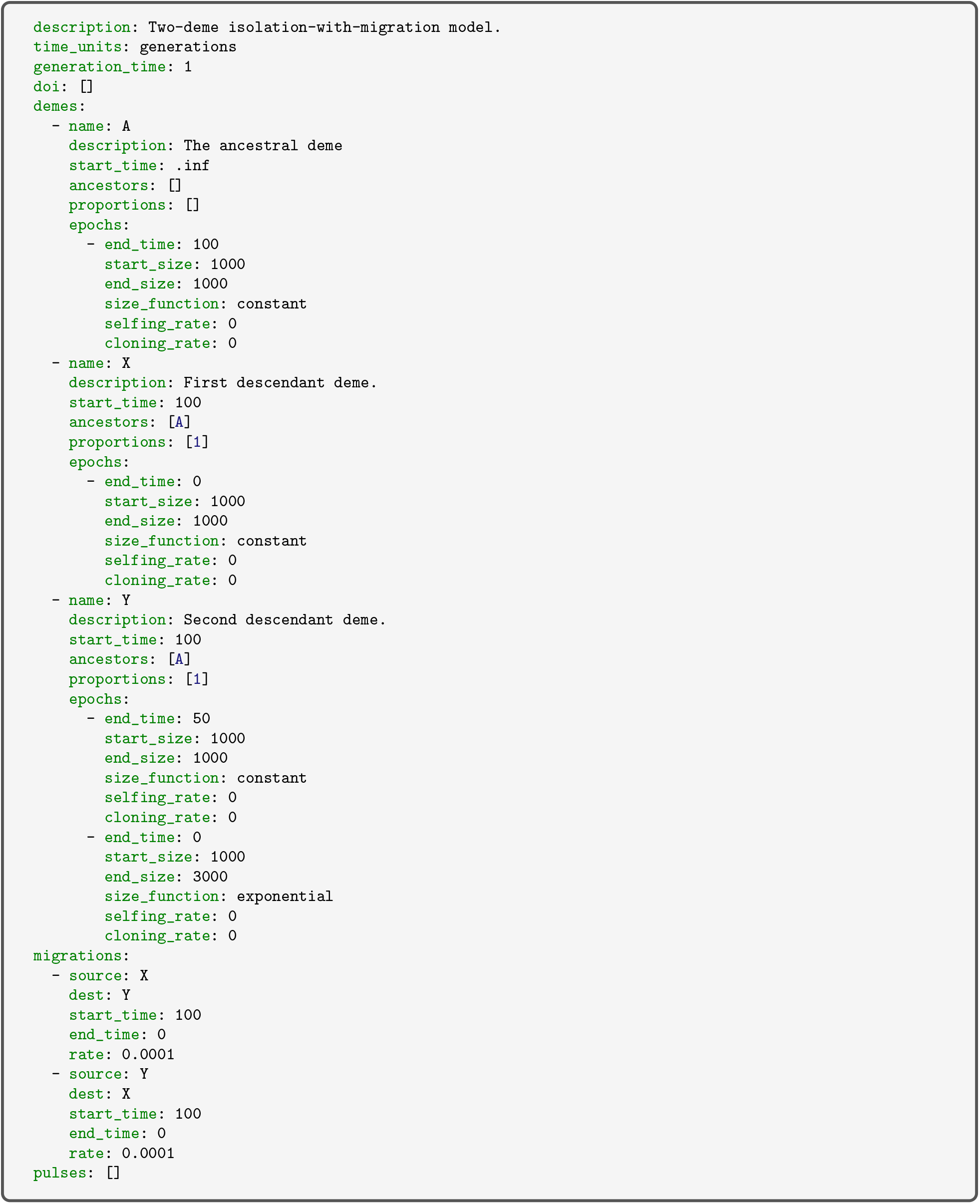
Isolation-with-migration example model from Figure 1 in Machine Data Model (MDM) form. The MDM form of the model is complete and explicit, but contains much redundant information that is omitted in the Human Data Model (HDM) form.

**Figure A2:**
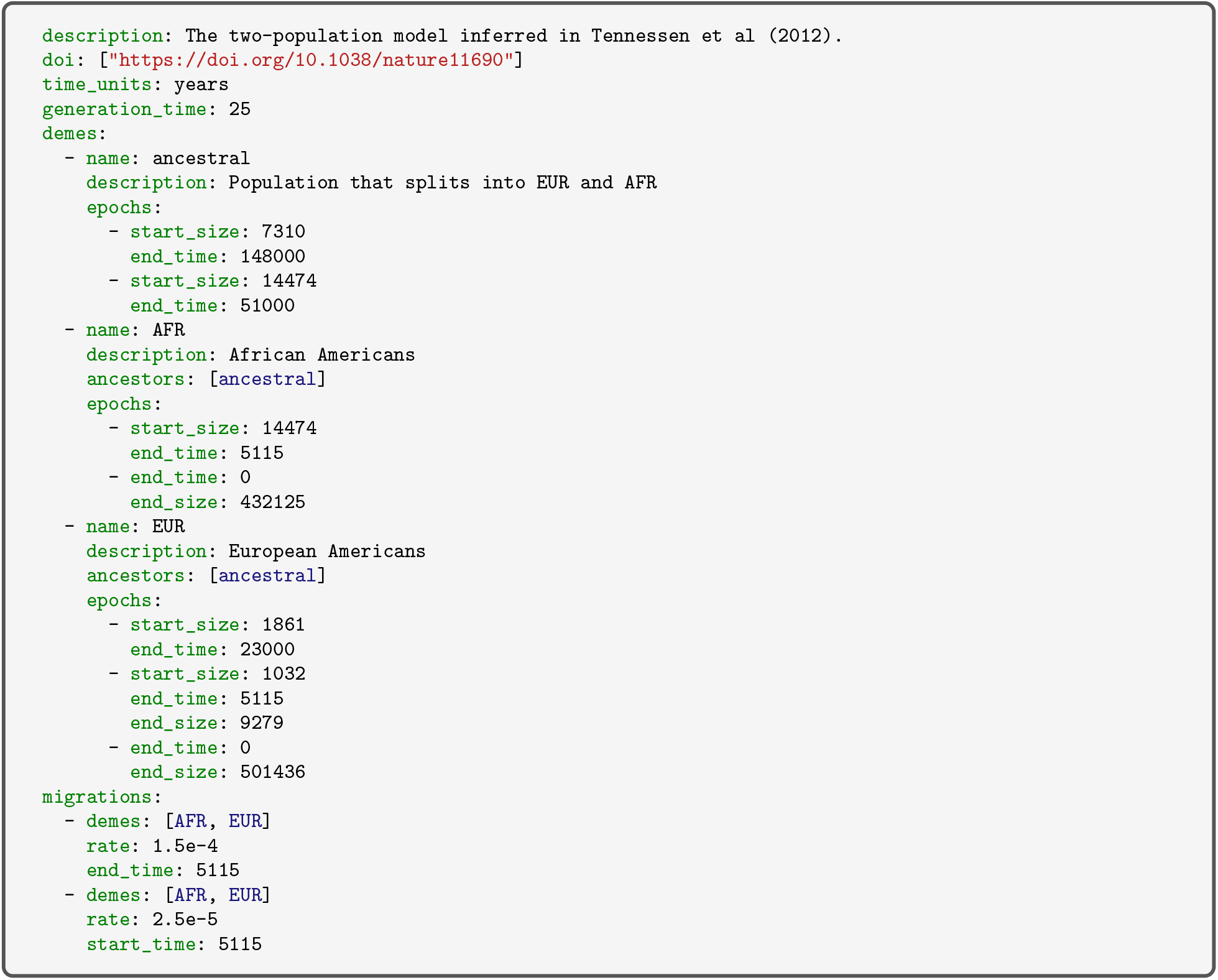
The Tennessen *et al*. (2012) two-population demographic model in Demes format. This model includes a single ancestral population that expands in size in the past, followed by divergence between AFR- and EUR-labeled populations. The two-population phase of the model includes multiple epochs of varying size, and rapid exponential growth over the past five thousand years in each population.

**Figure A3:**
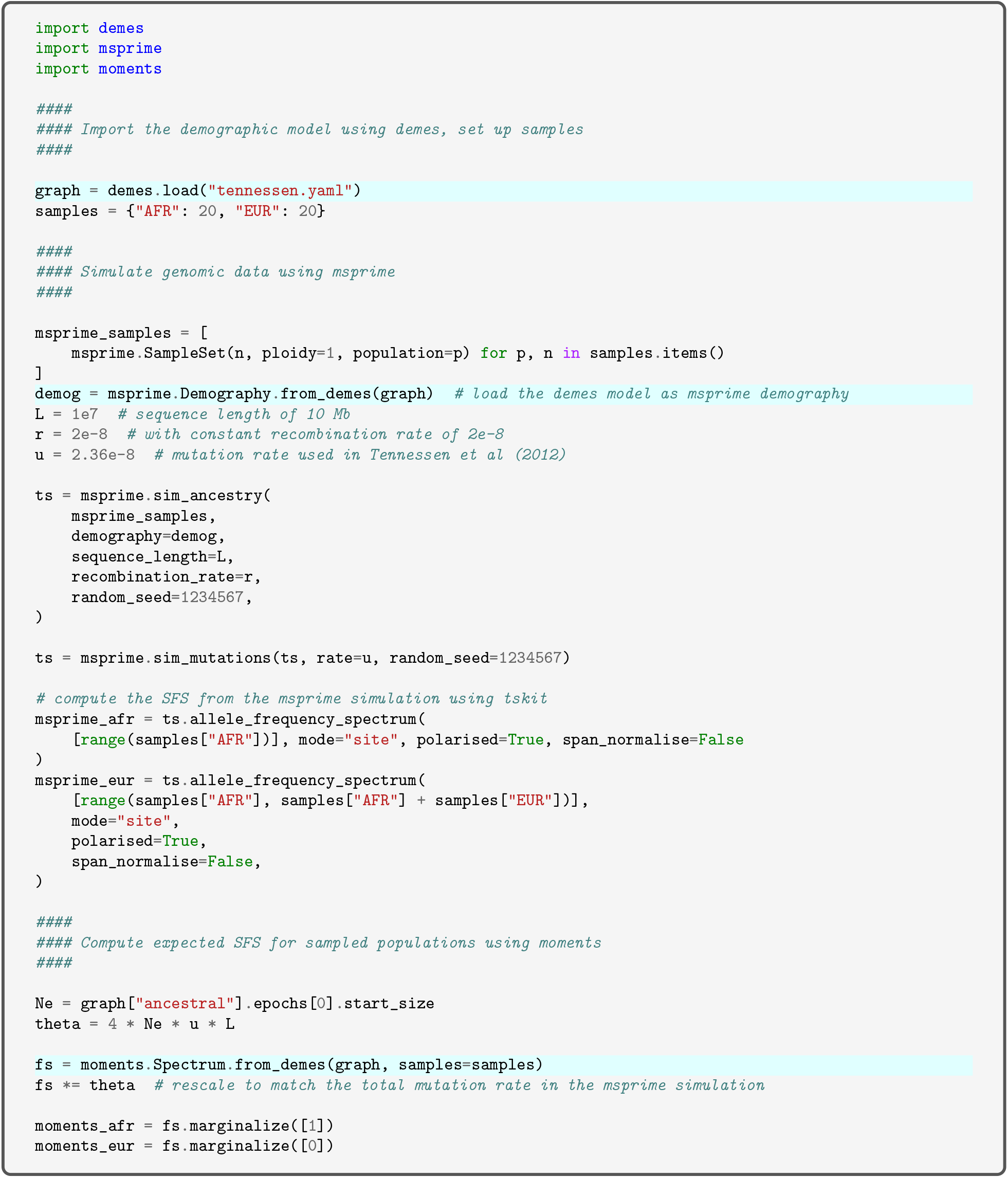
Simulation of SFS for the Tennessen model. We first load the demographic model using `demes` (as `graph`), which can then be used by `msprime` to create the demographic model used in `msprime.sim ancestry()`. The same loaded graph can also be passed to `moments` to compute the expected joint SFS. To compare the SFS in Figure 2, we marginalize the joint SFS to obtain the single-population SFS for both AFR and EUR populations. Lines interfacing `demes` and other software are highlighted.

## Notes

https://popsim-consortium.github.io/demes-spec-docs/

### Competing Interest Statement

The authors have declared no competing interest.

### Summary of Updates

Improvements to manuscript.

